# Transthyretin has conformation-selective proteolytic activity against α-synuclein

**DOI:** 10.1101/2023.08.10.552896

**Authors:** Zsuzsa Sárkány, Tiago Gião, Márcia Almeida Liz, Antoni Planas, Sandra Macedo-Ribeiro, Isabel Cardoso, Gemma Arsequell, Pedro M. Martins

## Abstract

Transthyretin (TTR) is a plasma protein known as a transporter of thyroxine and retinol but also can inhibit the formation of amyloid β-peptide (Aβ) fibrils and catalyze the proteolysis of apolipoprotein A-I and Aβ. Here, recombinant TTR is shown to have proteolytic activity against specific conformations of α-synuclein (aSyn), a protein that accumulates in intraneuronal inclusions characteristic of Parkinson’s disease (PD). Our discovery stemmed from the observation of a marked decrease in aSyn aggregation in the presence of submicromolar concentrations of TTR. Using integrated biophysical techniques, including advanced microscopy and mass spectrometry, we found that aSyn aggregation is prevented due to the proteolytic cleavage of free aSyn by TTR, in a reaction that is slower for S-glutathionylated TTR and faster for TTR preparations containing vestigial amounts of ∼70 kDa TTR oligomers (oTTR). Interestingly, this proteolysis-trigger effect is not unique to oTTR since it can also be induced by Teflon and glass surfaces independently of which TTR preparation is tested. Our results indicate that a change in aSyn conformation must precede the proteolysis step. The proteolysis of misfolded aSyn emerges as a possible TTR function with implications for the understanding of different neurodegenerative disorders.

## Introduction

Transthyretin (TTR) is present in the blood plasma and cerebrospinal fluid as a tetramer that transports thyroxine and the complex formed between retinol and the retinol-binding protein. Besides these transport functions, TTR is also able to proteolyze *in vitro* the natural substrates apolipoprotein AI (apoA-I) (1), neuropeptide Y (2), and amyloid β-peptide (Aβ) (3). Furthermore, TTR interacts with the oligomeric and fibrillar forms of Aβ acting as a potent inhibitor of Aβ aggregation and as a possible protector against neurodegeneration in Alzheimer’s disease (AD) (4–11). Different point mutations of TTR are related to Transthyretin amyloid polyneuropathy (ATTR-PN), a neurodegenerative disease characterized by the extracellular deposition of amyloid fibrils of TTR around peripheral nerves (12). Additionally, in non-hereditary TTR amyloidosis, amyloid fibrils of misfolded wild-type TTR accumulate in the heart tissue of elderly patients thus leading to severe cardiomyopathy (13).

The presynaptic protein α-synuclein (aSyn) has unclear physiological functions (14), is found mutated in some familial cases of Parkinson’s disease (PD), and is a major component of Lewy body inclusions characteristic of PD and dementia with Lewy bodies (15). The protein sequence of aSyn comprises a central non-amyloid-β component (NAC) region (residues 61−95) that is necessary for aSyn aggregation into the pathological filaments (16). α-Synuclein is a typical intrinsically disordered protein but, depending on the environment, can adopt several different conformational states (17). Within the N-terminal and NAC regions are seven highly-conserved imperfect 11-mer repeats that bear resemblance to sequence motifs found in ApoA-I and are believed to constitute amphipathic helical lipid-binding domains (18,19). Although the conformations of aSyn in-cell show structural disorder and are more compact than in buffer (20), the N-terminal and NAC regions are confirmed to fold as single or anti-parallel α-helices when bound to lipid membranes *in vitro* (19).

Interactions of aSyn with amyloid-forming proteins have been reported involving, for example, the most abundantly expressed extracellular protein, human serum albumin (21,22), the Aβ-peptide (23,24), and Tau (25,26). Protein chaperones, most notably heat shock proteins 70 and 90 (HSP70 and HSP90) and clusterin, modulate the structure, aggregation and toxic effects of different amyloidogenic proteins, including extracellular aSyn (27–29). Neuroserpin was recently added to the growing list of extracellular chaperones targeting different misfolding proteins (30). Importantly, although neuroserpin and TTR share a highly conserved 14-residue region and both are potent inhibitors of Aβ aggregation, only neuroserpin was found to efficiently inhibit aSyn aggregation: an excess of native (tetrameric) TTR had no evident effect on the amount of aSyn fibrils produced over time, while genetically engineered monomeric TTR showed limited inhibitory effects even when used at a molar excess of 50:1 to aSyn (30). Moreover, the spontaneous aggregation of aSyn seems unaffected by physiological concentrations of TTR (31). Simultaneously with the publication of these recent studies, we were observing a strong inhibition of aSyn aggregation by unmodified TTR but not by glutathionylated TTR. Intriguingly, the observed inhibition was even stronger if small amounts (<10%) of ∼70 kDa TTR oligomers (oTTR) were present in the unmodified, recombinant TTR preparations.

In the present work, evidence is provided that aSyn is cleaved by TTR into aggregation-incompetent fragments. We integrate multiple biophysical approaches, including fluorescence spectroscopy, electron microscopy, gel electrophoresis and mass spectrometry to show that the conformation of aSyn must be changed before TTR-mediated proteolysis occurs. In addition to explaining the effects of TTR on aSyn aggregation, the discovery of a new protease with conformation-selective activity against aSyn is a fundamental breakthrough that opens new possibilities for investigating PD pathogenesis and treatment.

## Results

### α-Synuclein aggregation is affected by the concentration and oligomeric composition of TTR

To study the effect of TTR on the aggregation kinetics of aSyn, Thioflavin T (ThT) fluorescence was monitored over ∼180 h during the aggregation of 2 μM aSyn in the presence of 0.2 μM pre-formed fibrils (PFFs) of aSyn and serially diluted concentrations of TTR (Figs. S1a and S1b). Titrations starting at 20 μM TTR yield half-maximal inhibitory concentrations (IC_50_) of 6 nM TTR (Fig. 1a). Yet, when TTR is serially diluted from a 2.5 μM solution, the inhibitory effect of TTR becomes weaker (IC_50_=64 nM TTR). No disaggregation effect is identified by adding TTR to aSyn PFFs (data not shown). Size exclusion chromatography (SEC) and SDS polyacrylamide gel electrophoresis (PAGE) analyses revealed that 50 μM TTR samples contained ∼93% tetrameric TTR (tTTR) and ∼7% of TTR oligomers (oTTR) with an estimated molecular weight of ∼70 kDa as shown by SEC (Fig. 1b). The fraction of oTTR decreases to ∼4% when 5 μM TTR solutions are analysed (Fig. S2a) suggesting that a concentration-dependent equilibrium is established between tTTR and oTTR. From these results, we hypothesized that oTTR species were, at least in part, responsible for the observed inhibition effect on aSyn aggregation: departing from more concentrated TTR solutions would assure higher oTTR/tTTR ratios and, consequently, stronger inhibition. To discriminate the contributions of oTTR and tTTR for the observed inhibition, the corresponding SEC fractions were collected, concentrated and tested independently during aSyn aggregation in new dose-response studies (Figs. S1c and S1d). When departing from concentrated oTTR solutions, the estimated IC_50_ value (0.147 μM TTR) is still much higher than the obtained for the mixture tTTR and oTTR (6 nM TTR) under equivalent conditions (Fig. 1c). The tetrameric tTTR produced no significant inhibition of aSyn aggregation as identified both by ThT fluorescence (Fig. 1c) and TEM (Fig. 1d). A key role of oTTR is, therefore, confirmed, yet the inhibitory effect elicited by the sample containing tTTR and oTTR is not exclusively due to oTTR.

**Figure 1.**
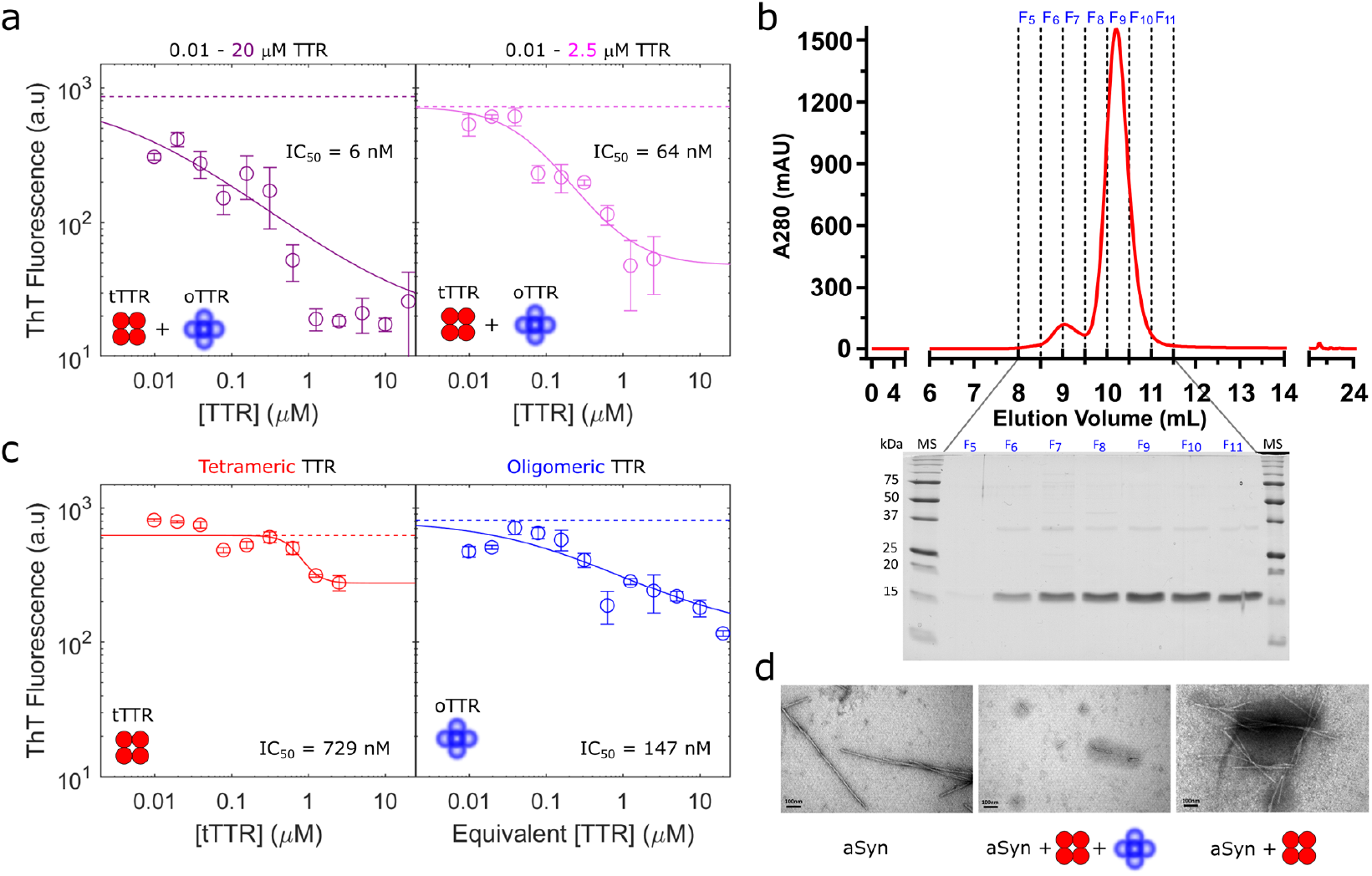
α-Synuclein aggregation is affected by the concentration and oligomeric composition of TTR. (a) Mean (symbols) and standard deviation (error bars) of end-point ThT fluorescence values measured after the seeded aggregation of 2 μM aSyn in the presence of serially diluted 20 μM (left) and 2.5 μM (right) TTR. Solid lines: numerical fittings by a concentration-response curve. Dashed lines: mean endpoint fluorescence measured during control experiments with no TTR. (b) Top: SEC chromatogram of 50 μM TTR; F5 to F11 indicate different collected fractions. Bottom: 15% SDS-PAGE analysis of the SEC fractions collected after injection of 50 μM TTR. (c) Influence of tTTR (left) and oTTR (right) on the end-point ThT fluorescence. In the case of oTTR (right), an equivalent TTR concentration is estimated from the number of times the eluted sample is concentrated. Symbols, error bars and lines as in (a). (d) Transmission Electron Microscopy (TEM) images of negatively stained aliquots collected at the end-point of the seeded aggregation of 2 μM aSyn in the absence (left) and presence of 2.5 μM TTR samples containing oTTR (centre) and oTTR-free (right); scale bars: 100 nm. No nucleation or elongation of aSyn fibrils is detected when oTTR is present (centre).

### TTR-mediated proteolysis of α-synuclein

Polyacrylamide gel electrophoresis (PAGE) followed by western blotting immunoassays were used to further characterize the species present in the TTR sample (Fig. 2a) and the possible aSyn aggregates generated in the presence of TTR without aSyn PFFs (Fig. 2b). It could be the case that oTTR was inducing the off-pathway aggregation of aSyn and the concomitant formation of fewer ThT-positive assemblies during seeded aggregation of aSyn (32). Native-PAGE analysis of the TTR sample (Fig. 2a, left) confirmed the occurrence of oTTR species, which, curiously, immunostained with the anti-TTR antibody more intensely than tTTR (Fig. 2a, left). In support of the view that oTTR coexisted with tTTR in a dynamic equilibrium, the relative densities of immunostained oTTR and tTTR seem to vary with the total amount of protein loaded into the gel (Fig. 2a, left). The western blot analysis of aliquots of aSyn incubated with the mixture tTTR/oTTR shows the disappearance of the aSyn band and the appearance of a lower band corresponding to proteolysis, without the simultaneous formation of aSyn oligomers (Fig. 2b). Similarly, the formation of ASyn or TTR aggregates is not detected by SEC (Figure S2b) and DLS (Figure S2c). The analysis of the rates of aSyn decay indicates a catalytic efficiency (k_cat_/K_M_) of 197 s^−1^M^−1^ that is 67× higher than the value estimated using the Aβ (1–42) peptide as substrate (Fig. S3).

**Figure 2.**
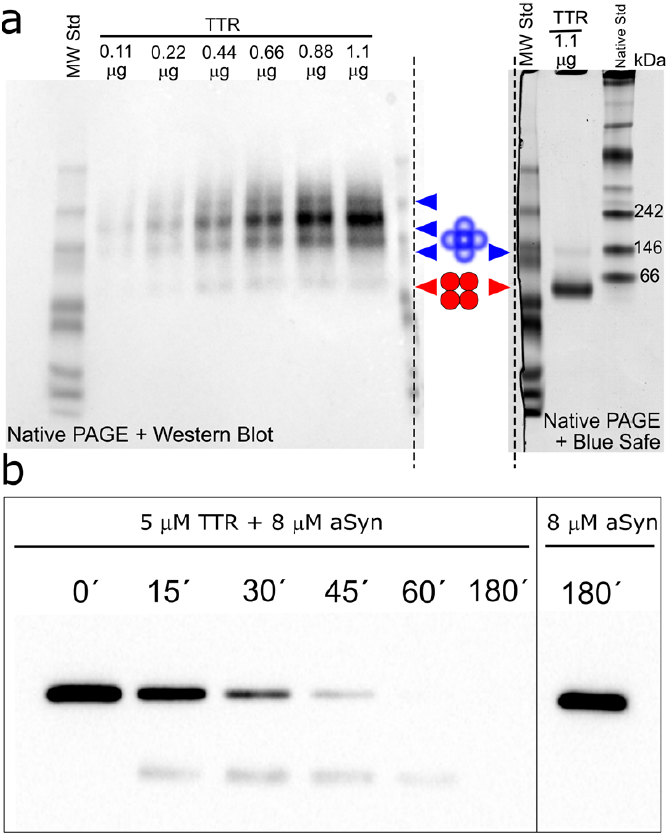
Immunoblotting of TTR and TTR/aSyn samples samples. (a) Native-PAGE of the TTR sample analysed by western blot (anti-TTR antibody) (left), and stained with Blue Safe protein stain (right). The gel was vertically cut as indicated by the dashed lines. Blue and red arrows indicate oTTR and tTTR, respectively. (b) SDS-PAGE western blot analysis of aliquots of 8 μM aSyn incubated with 5 μM TTR indicates aSyn proteolysis by TTR. The aliquots were immunoblotted with the anti-aSyn antibody.

We then performed Matrix-Assisted Laser Desorption/Ionization coupled to Time-of-Flight Mass Spectrometry (MALDI-ToF-MS) analysis to follow aSyn proteolysis by TTR (Fig. S4). From time points 0 min to 15 min, the emergence of two major peaks of 5394.2 Da and 6544.7 Da was observed. These fragments become more evident at time point 30 min and remain visible at time points 45 min and 60 min together with several other new fragments. At time point 180 min, the peaks are too numerous to be accurately analysed. To unambiguously identify the aSyn fragments in the proteolysis, a top down MS approach using nanoLiquid Chromatography coupled to high-resolution Mass Spectrometry (nanoLC-MS/MS) was employed at time points 15, 30 and 45 min (Fig. 3). A combination of different MS acquisition methods and fragmentation techniques was used to obtain sequence coverage of as many proteoforms as possible. We found five N-terminal proteolytic fragments of aSyn covering the following regions: 1-53, 1-64, 1-72, 1-79 and 1-92 and five C-terminal proteolytic fragments of aSyn covering the following regions: 54-140, 73-140, 80-140, 90-140 and 91-140.

**Figure 3.**
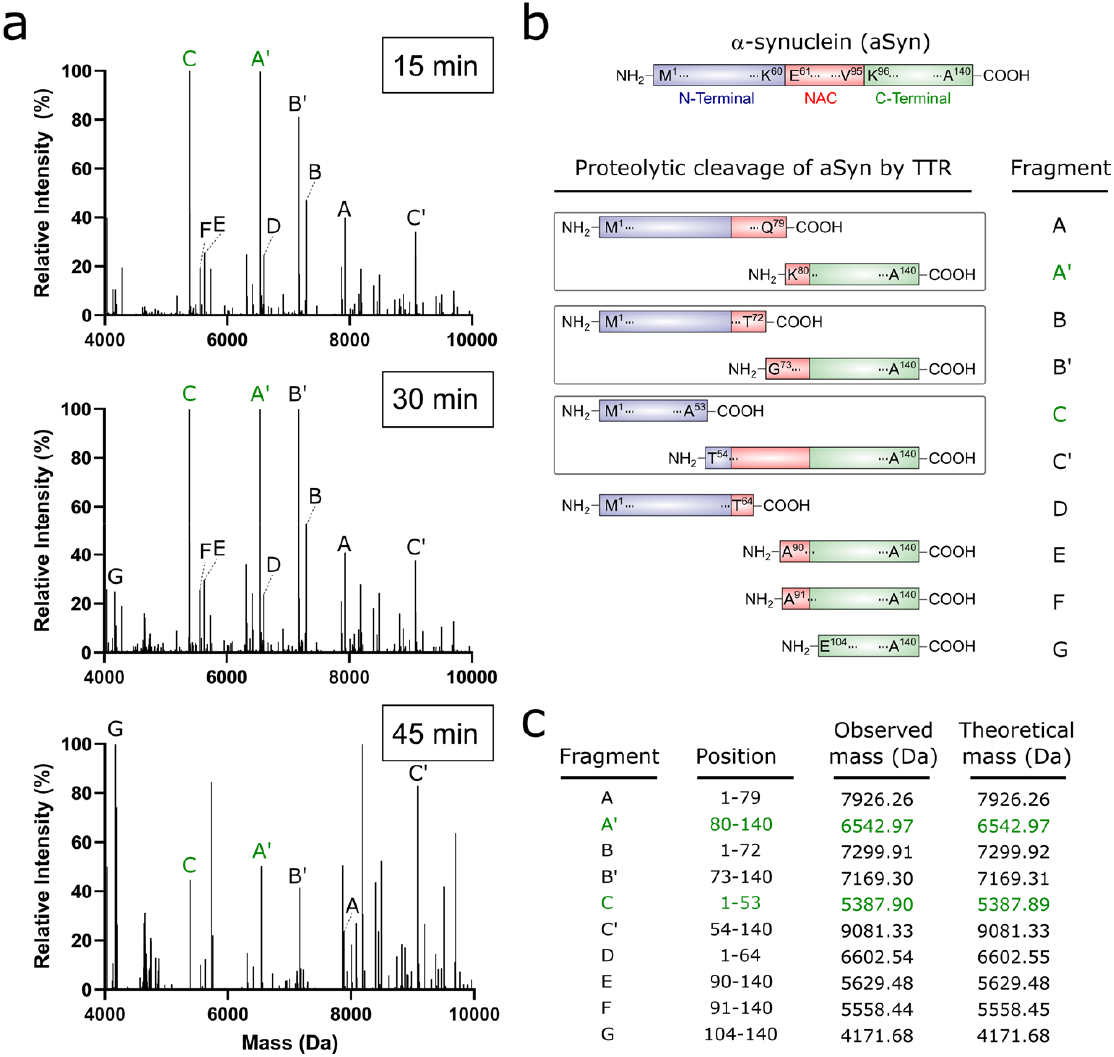
Determination of main proteoforms from aSyn proteolysis by TTR. (a) Xtract (36) deconvoluted mass spectra from 4000 Da to 10000 Da of 8 μM aSyn incubated with 5 μM TTR over periods of (top) 15, (middle) 30, and (bottom) 45 min. Letters next to the peaks indicate different aSyn fragments. Green letters indicate abundant fragments. (b) Schematic illustration of full-length aSyn and the cleavage sites for TTR in aSyn. Summary of the start/stop residue positions, the observed mass, and the theoretical mass for each proteolysis proteoform detected. The supporting nanoLC-MS/MS data are provided in SI Appendix (Table S1).

We confirmed the existence of two abundant fragments corresponding to 1−53 aSyn and 80−140 aSyn with masses 5386.4 Da and 6543.0 Da, respectively (marked green in Fig. 3). The complementary fragments 54−140 aSyn and 1−79 aSyn were also identified thus suggesting two major cleavage sites for TTR in aSyn at Ala^53^/Thr^54^ and Gln^79^/Lys^80^. Notably among the amino acids involved, Ala^53^ and its mutation to Thr is the first hereditary mutation of aSyn identified in families with autosomal dominant early-onset PD (33). An additional cleavage site at Thr^72^/Gly^73^ is suggested by the identification of fragments 1−72 aSyn and 73−140 aSyn. Altogether, the mass spectrometry evidence rationalizes why aSyn cleavage by TTR generates aggregation-incompetent fragments because the cleavage sites are located within the central NAC (residues 61−95) and pre-NAC (residues 45–57) regions that are essential for aSyn aggregation (34,35).

### α-Synuclein is cleaved by tTTR in the absence of oTTR and in a conformation-selective way

To elucidate the roles of the TTR species in the aSyn proteolysis, immunoblotting was used to quantify aSyn after incubation with tTTR, oTTR or a mixture tTTR:oTTR (10:1) prepared after isolating each species by SEC), independently. Proteolytic activity is identified for the tTTR:oTTR preparation but not for isolated tTTR or oTTR (Fig. 4a, top). The aSyn band strongly decreases upon long incubation with tTTR+oTTR but, in this case, the proteolysis fragments are barely detected (as they were in Fig 2a up to 1h incubations) suggesting further degradation. Since three tTTR substrates have been reported so far (1–3), we hypothesized that the proteolytic cleavage of aSyn is also assured by tTTR although prior action of oTTR is required to turn aSyn susceptible to proteolytic attack. Conformation changes of aSyn could be induced either by specific aSyn-oTTR complexes or by other types of interaction such as those established, for example, with polytetrafluoroethylene (PTFE or Teflon) surfaces (37,38). The second of these possibilities was confirmed by immunoblotting experiments showing that the proteolytic activity of isolated tTTR is displayed under quiescent conditions in the presence of a Teflon surface (Fig. 4a, bottom). Only 31% aSyn remained in solution after 20 h incubation with tTTR in the presence of a Teflon-coated stir bar, whereas no evident decrease of the aSyn band intensity is observed in the control experiments without tTTR and/or Teflon bars. Incidentally, the Teflon surface also appears to promote the spontaneous degradation of aSyn to a very small extent, only evidenced by the faint smeared region down the length of the corresponding lane (Fig. 4a, bottom). In an attempt to identify aSyn conformers selectively targeted by tTTR, high-sensitivity fluorescent immunoblotting was used to analyse the result of aSyn incubation with and without the presence of aggregation-interfering agents. A smeared region above the aSyn band does appear after 20 h incubation in the presence of Teflon surfaces, but is not seen in the control experiment nor in the presence of isolated oTTR (Fig. 4b). In the presence of oTTR, the intensity of the aSyn band is more fainted as a possible consequence of the proteolytic activity of tTTR that is either dissociated from oTTR or co-eluted in the SEC fraction of oTTR.

**Figure 4.**
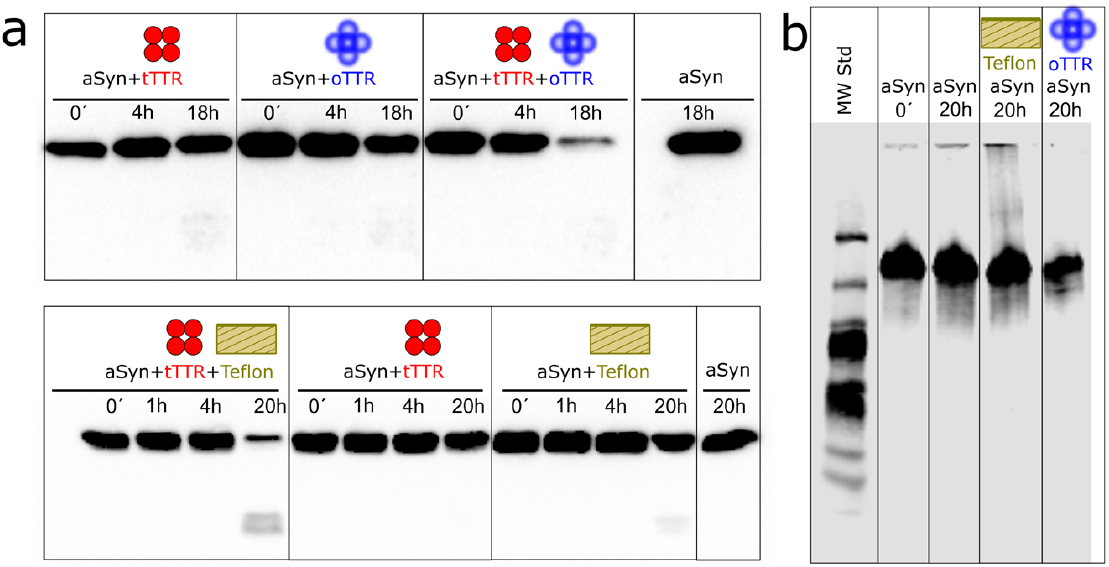
Identification of oTTR and Teflon surfaces as aSyn-conformation modifiers under quiescent conditions. (a) Top: Western blot analysis of 8 μM aSyn aliquots incubated in LoBind polypropylene test tubes with 5 μM tTTR (left series), 0.5 μM oTTR (centre series) and the mixture 5 μM tTTR:0.5 μM oTTR (right series) over the indicated time intervals. Far-right lane: negative control corresponding to 18 h incubation of 8 μM aSyn. Tetrameric TTR alone does not proteolyze aSyn effectively. Bottom: Western blot analysis of 8 μM aSyn aliquots incubated in polypropylene test tubes with 5 μM tTTR and a Teflon bar (left series), 5 μM tTTR without a Teflon bar (centre series) and no tTTR and a Teflon bar (right series). Far-right lane: negative control corresponding to 20 h incubation of 8 μM aSyn. Similar to oTTR, Teflon surfaces facilitate aSyn proteolysis by tTTR. (b) Native-PAGE, fluorescent western blot analysis of 8 μM aSyn solutions freshly prepared (second lane) or incubated over 20 h in the absence (third lane) or presence of a Teflon bar (fourth lane) or 0.5 μM oTTR (fifth lane). An anti-aSyn antibody was used in (a) and (b). The Teflon bar originates different conformational states of aSyn (upper smeared band); similar bands are not identified in the presence of only oTTR most likely due to aSyn proteolysis.

To test if the hydrophobic nature of Teflon surfaces was responsible for the conformation changes in aSyn, the effect of a hydrophilic surface (glass) on aSyn proteolysis by tTTR was investigated under quiescent conditions (Fig. 5a). Glass beads with a total surface area equivalent to the Teflon bar were added to aSyn/tTTR solutions and then soluble aSyn was monitored by western blotting (Fig. 5a). The main products of aSyn proteolysis by TTR were identified using MALDI-ToF-MS (Fig. S5). Since glass surfaces accelerate aSyn cleavage even more than Teflon surfaces, we conclude that the formation of proteolysis-prone aSyn conformers is also possible in the presence of hydrophilic surfaces. Control experiments run in the absence of TTR show that the presence of the solid surfaces alone does not provoke a visible change in the amount of intact aSyn at the end of 26 h incubation (Fig. S6).

**Figure 5.**
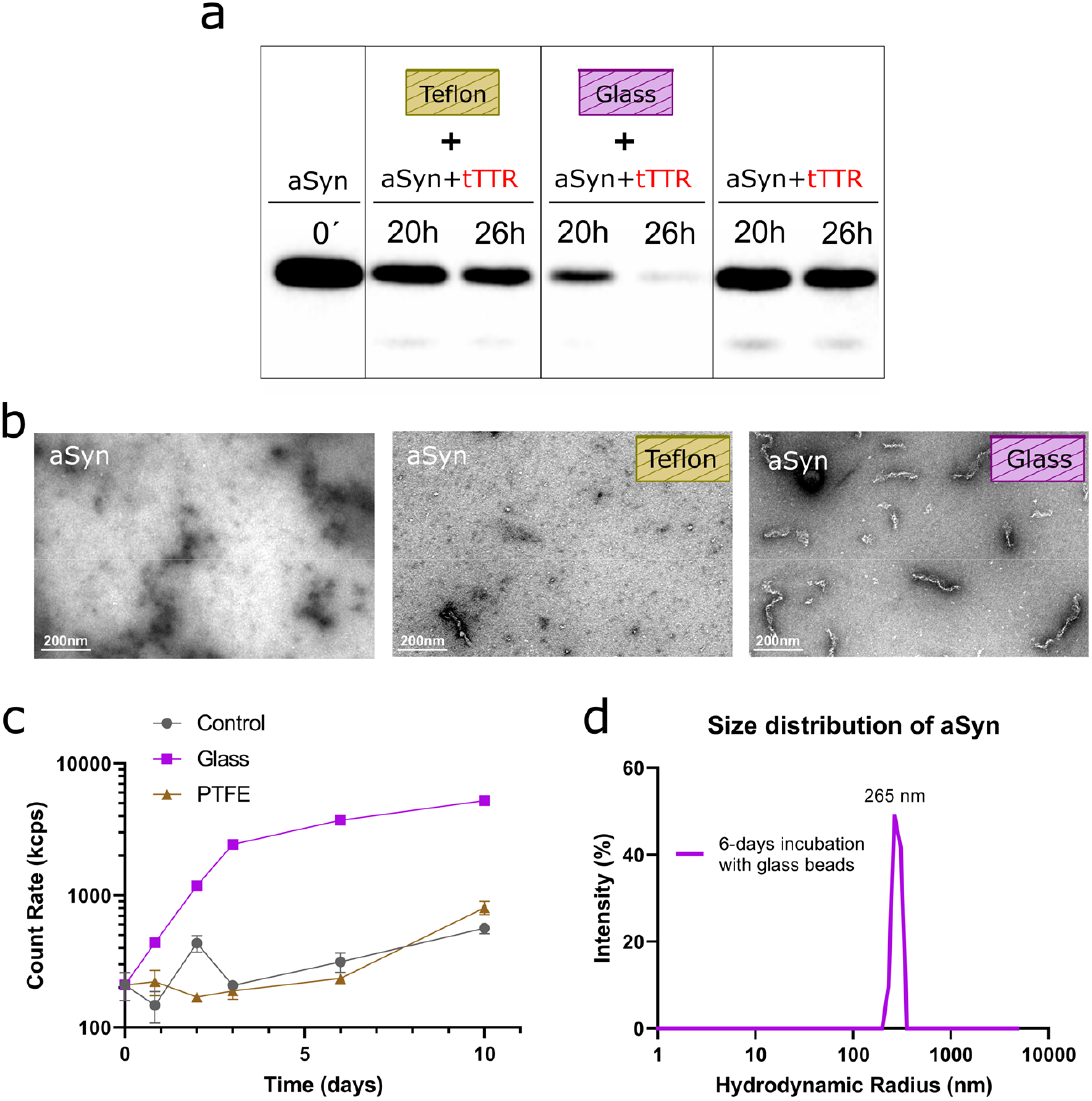
Glass surfaces are stronger aSyn-conformation modifier than Teflon surfaces. (a) Western blot analysis of 8 μM aSyn aliquots incubated in LoBind polypropylene test tubes with 5 μM tTTR in the presence of Teflon (second and third lanes) and glass (fourth and fifth lanes) surfaces over the indicated time intervals. First lane and far right lanes: controls in the absence of Teflon or glass surfaces. (b) TEM images of negatively stained aliquots collected after 12 days incubation of 8 μM aSyn in the absence (left) and presence of Teflon (middle) or glass (right) surfaces; scale bars: 200 nm. (c) Increase of light scattering intensity during the incubation of 8 μM aSyn in the presence of the different surfaces (log-linear scale). (d) DLS-intensity size distributions of 8 μM aSyn samples collected after 6 days of incubation in the presence of glass beads. (b to d) TTR-free experiments.

We then questioned what was the effect of Teflon and glass surfaces on aSyn aggregation in the absence of TTR and considering longer periods than 1 day. For the low aSyn concentrations used in our assays (8 μM), no measurable increase in ThT fluorescence was identified at the end of 12 days incubation (results not shown). However, TEM observations of aSyn samples incubated in the presence of glass surfaces revealed the presence of abundant fibrillar species of ∼200 nm length (Fig. 5b). Compared with mature aSyn fibrils generated in the presence of PFFs (Fig. 1d, left), the species now identified are shorter and show curvilinear rather than straight shapes. Similar protofibrils but much less abundant could be observed in the samples incubated in the presence of the Teflon bar; no aggregates were evident when aSyn was incubated in the absence of Teflon or glass surfaces (Fig. 5b). In line with these observations, the light scattering intensity increases more pronouncedly over time when in the presence of glass beads (Fig. 5c). As a result, reproducible size distribution measurements were only possible using the samples incubated with glass beads. A dominant peak centred in 265 nm hydrodynamic radius is identified (Fig. 5d) in accordance with the analysis of TEM micrographs (Fig. 5b). Taken together, these results suggest that proteolysis-prone conformers of aSyn are readily produced in the presence of glass surfaces and, to a less extent, Teflon surfaces. Unless they are proteolyzed by TTR, these aSyn conformers will tend to aggregate into short, curvilinear fibrils. In conclusion, we propose a mechanism in which, first, new aSyn conformers are generated from interactions with certain surfaces or oTTR, and, second, the modified aSyn molecule is proteolyzed by tTTR.

### Modulation of α-Synuclein proteolysis by the oligomeric content and S-glutathionylation of TTR

S-Glutathionylation of the Cys-10 residue of TTR was previously shown to abolish the proteolytic activity of TTR without affecting its functions of thyroxine and retinol transport (2,39). To investigate whether this post-translational modification also affects aSyn proteolysis, recombinant TTR preparations from different sources and containing different levels of S-glutathionylated (S-GSH) and oligomeric protein were tested. Specifically, we compared aSyn proteolysis by (i) the wt TTR produced using our first purification protocol (TTR_1st_) with aSyn proteolysis by (ii) the highly amyloidogenic Y78F TTR mutant purified by the same protocol (Y78F_1st_) and (iii) the wt TTR purified according to a second protocol (TTR_2nd_) that yields less oligomeric TTR (Fig. 6a). The first purification protocol includes a step of fractional ammonium sulfate precipitation (40), whereas the second protocol includes a step of preparative gel electrophoresis (3). By employing MALDI-ToF-MS, we further detected the unmodified and S-GSH forms of TTR (40,41), and the corresponding peak intensities were used to characterize unmodified to S-GSH TTR ratios (Figs. 6b and S7). Besides containing less oligomeric species, the TTR_2nd_ source has higher amounts of S-glutathionylated protein (S-GSH-TTR_2nd_).

**Figure 6.**
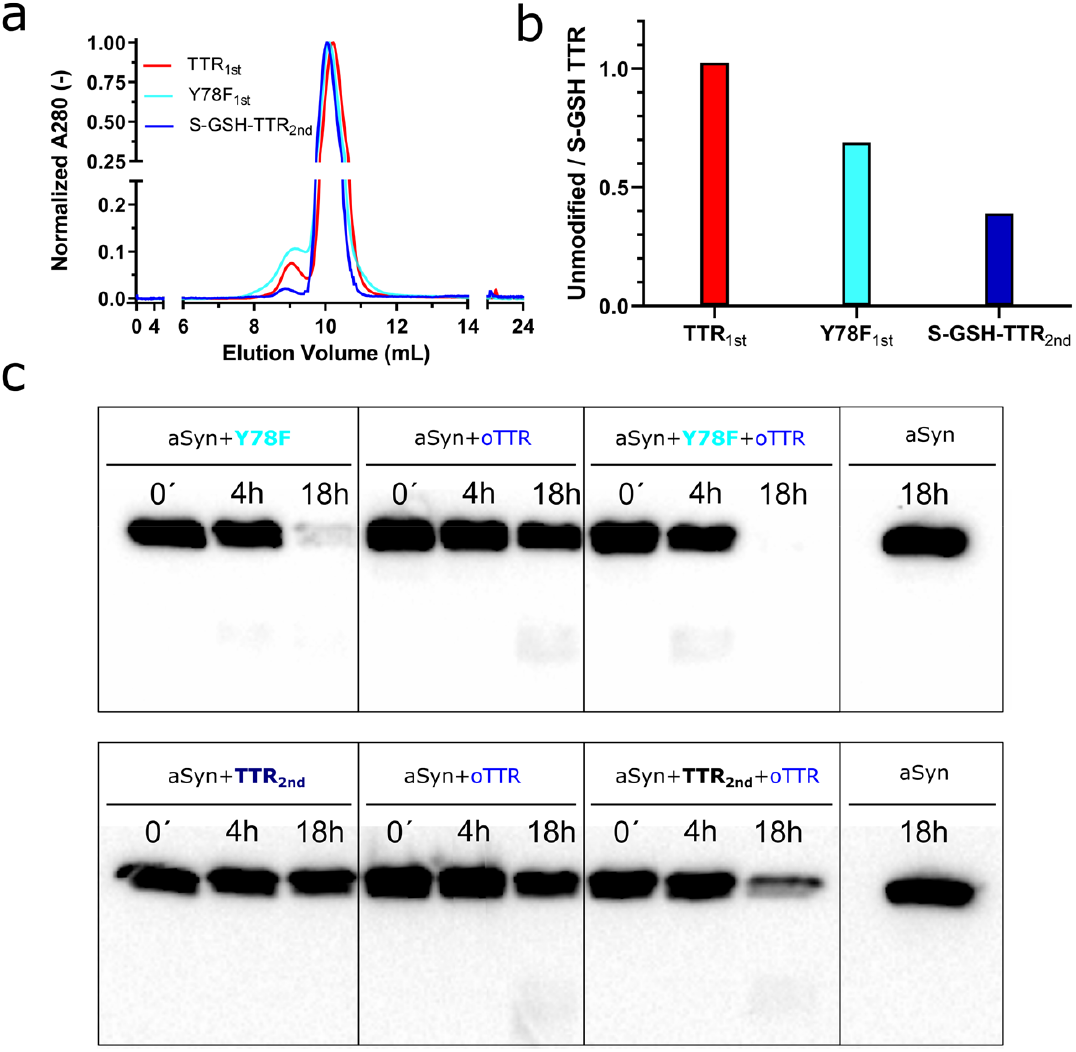
The effects of S-glutathionylation and oligomeric content of TTR on its proteolytic activity. (a) Normalized SEC chromatograms of (red) TTR_1st_, (light blue) Y78F_1st_ and (dark blue) S-GSH-TTR_2nd_ sources. (b) Ratios of unmodified to S-GSH TTR estimated from MALDI-ToF-MS spectra (Fig. S7) in TTR samples of different sources. (c) Western blot analysis of aSyn proteolysis upon incubation with (top) Y78F_1st_ and (bottom) S-GSH-TTR_2nd_ over the indicated time intervals. Top: incubation of 8 μM aSyn with 5 μM Y78F_1st_ (left series), 0.5 μM oTTR (centre series) and the mixture 5 μM Y78F_1st_:0.5 μM oTTR (right series). Bottom: incubation of 8 μM aSyn with 5 μM S-GSH-TTR_2nd_ (left series), 0.5 μM oTTR (centre series), and the mixture 5 μM S-GSH-TTR_2nd_:0.5 μM oTTR (right series). Far-right lanes: negative controls corresponding to 18 h incubation of 8 μM aSyn. The proteolysis of aSyn by S-GSH TTR requires the presence of aSyn-conformation modifiers such as oTTR.

An inverse correlation was confirmed between the S-GSH level and the proteolytic activity of TTR: full proteolysis of aSyn by TTR_1st_ is completed in ∼1 h (Fig. 2b), Y78F_1st_ requires >18 h to achieve it, while S-GSH-TTR_2nd_ appears to have no proteolytic activity (Fig. 6c, top). No correlation between activity and oligomeric content of TTR was found because Y78F_1st_ has more TTR oligomers and less catalytic activity than TTR_1st_. It should, however, be considered that the chemical nature of the oligomers present in each of these samples is different. Moreover, it remains to be demonstrated whether S-GSH-TTR_2nd_ is active in the presence of modifiers of aSyn conformation. To address these open questions, oTTR extracted from TTR_1st_ was used to prepare 1:10 mixtures of oTTR:Y78F_1st_ and oTTR:S-GSH-TTR_2nd_. A marked increase in proteolytic activity was observed in both cases. Quantitative immunoblotting indicates that the amount of proteolyzed aSyn at the end of 18 h from 91% to 100% and increased from ∼0% to 64% when oTTR is present in the Y78F_1st_ and S-GSH-TTR_2nd_ preparations, respectively (Fig. 6c, bottom). The proteolysis-promoting effect of oTTR on S-GSH-TTR_2nd_ is comparable to the 72% increase observed under identical conditions for tTTR preparations extracted from TTR_1st_ (Fig. 4a). This indicates that the proteolytic activity of S-GSH TTR can be displayed in the presence of modifiers of aSyn conformation. To investigate whether the effect of S-glutathionylation is dependent on the protein production protocol, a different batch of TTR_2nd_ having unmodified Cys-10 (unmod-TTR_2nd_) is shown to be more proteolytically active against aSyn (Fig. S8). Overall, these results demonstrate that aSyn proteolysis by TTR is exacerbated in the presence of modifiers of aSyn conformation and inhibited by Cys-10 S-glutathionylation.

## Discussion

The causes for the apparent inhibition of aSyn aggregation by TTR are here traced back to an unexpected proteolytic activity of tTTR against specific conformations of aSyn. A relationship was initially established between aSyn amyloid production and the relative amounts of tetrameric and oligomeric TTR in solution. A population of ∼70 kDa oTTR, which coexists with tTTR in a dynamic equilibrium, is found to drastically affect the seeded aggregation of aSyn (Fig. 1). The absence or presence of ∼7% oTTR in the TTR preparation determines a change in the inhibitory effect from nearly undetectable to very strong (IC_50_<10 nM). Crucially, a similar dependence on the oTTR content is identified for the rates of exhaustion of free aSyn incubated without aSyn PFFs (Figs. 2a, 2b and 4a). The correspondence between the two effects provoked by oTTR is elucidated using MALDI-ToF-MS and nano-LC-MS/MS (top-down approach) which show that the concentration of free aSyn decreases due to aSyn proteolysis into aggregation-incompetent aSyn proteoforms. The proteolysis-facilitating effect of oTTR is not specific to TTR oligomers since it is also induced in the presence of Teflon and glass surfaces (Fig. 7) that are known to promote aSyn aggregation (37,38). α-Synuclein incubated with TTR without conformation modifiers will not be proteolyzed unless the TTR sample has very low levels of S-glutathionylation. This explains why the effect of TTR on the integrity (and aggregation properties) of aSyn was not detected in earlier studies of protein-protein interaction (30,31).

**Figure 7.**
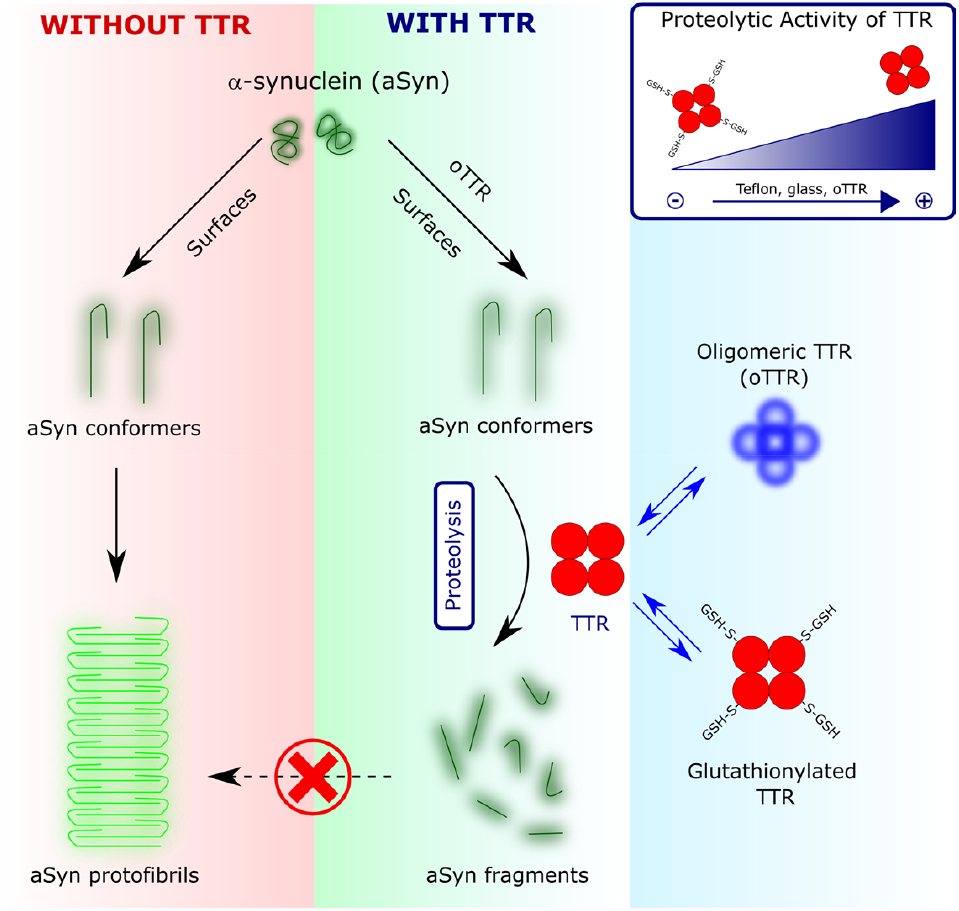
Schematic illustration of the conformation-selective proteolytic activity of TTR against aSyn. α-Synuclein cleavage by tTTR is accelerated in the presence of modifiers of aSyn conformation (such as Teflon, glass or oTTR) and modulated by the S-glutathionylation of tTTR. Unless the modified conformation of aSyn is proteolytically cleaved by TTR, it will aggregate into protofibrillar species.

Our findings raise important questions about the roles of TTR and aSyn in health and disease, viz.:

- Is the neuroprotective role of TTR in AD also verified in PD?
- Is the proteolysis of misfolded aSyn a physiological function of TTR?
- Can the proteolytic activity of TTR be regulated by post-translational modifications such as S-glutathionylation?
- Are there other proteases than TTR with the hidden ability to selectively target misfolded proteins other than aSyn?

To address these questions, it is worth reviewing the scarce evidence available showing the involvement of TTR in PD (42). In 1990, a 1.9-fold increase in the levels of cerebrospinal fluid (CSF) TTR was reported for PD patients who underwent adrenal medullary autotransplantation, an effect that was explained by the possible activation of the choroid plexus where CSF TTR is mainly produced (43). In 2007, a 2.4-fold increase in CSF TTR concentration was reported for a rat model recapitulating the nigrostriatal dopaminergic degeneration characteristic of PD. Interestingly, two works dated back to 2012 show, in one case, increased levels of monomeric and oligomeric aSyn in the saliva of symptomatic ATTR-PN patients; liver transplantation (where plasma TTR is mainly produced) resulted in aSyn levels in saliva indistinguishable from healthy and asymptomatic controls (44). In the other study, “(misfolded) aSyn” is hypothesized as “a target protein of TTR” because the CSF TTR levels in non-demented PD patients were significantly higher compared to controls, while demented patients with Lewy body disorders had relatively low CSF TTR levels (45). As concluded by the authors (45), aSyn pathology alone goes along with increased TTR expression but this effect was counteracted if Aβ pathology (and dementia) was also present (46).

Conversely, several lines of evidence point to TTR as neuroprotective (10,47), as well as a candidate biomarker (48,49) in AD. In addition to the significant negative correlation between CSF TTR levels and AD severity (46), a reduction of Aβ toxicity in hippocampal neurons is observed depending on the proteolytic activity of TTR (50). The discovery that TTR also proteolyzes aggregation-prone aSyn conformers and does it more efficiently (k_cat_/K_M_=197 s^−1^M^−1^) than during Aβ(1-42) cleavage (k_cat_/K_M_ =2.9 s^−1^M^−1^) might suggest a neuroprotective role of TTR in PD, in which case the increased expression of TTR would be a response to higher levels of misfolded aSyn.

α-Synuclein is predominantly presynaptic but is also found in the CSF, brain interstitial fluid and plasma. Clearance of extracellular aSyn is assured by proteases such as matrix metalloproteinases (MMPs) or through uptake into cells and subsequent degradation (51). PD pathogenesis and aSyn spreading to different regions of the brain have been associated with impaired anti-oxidative mechanisms required for the activation of MMPs targeting extracellular aSyn (51,52). Free and aggregated forms of aSyn are efficiently broken by MMPs such as MMP-2 and MMP-3 along multiple cleavage sites located outside and inside the NAC region of aSyn (35,52). Similar to MMPs, TTR has metalloproteinase activity whose inhibition by EDTA or 1,10-phenanthroline can be reverted by zinc and manganese addition (39). One of the major cleavage sites for TTR in aSyn − Gln^79^/Lys^80^ − is a common target to MMP3 (53,54). However, unlike what happens during the proteolysis of aSyn by MMP3, the multiple fragments generated in the presence of TTR are not aggregation-prone and do not include 79−140 aSyn, whose pyroglutamate modification promotes oligomer formation and neurotoxicity in human synucleinopathies (53). TTR also differs from MMPs, plasmin or neurosin because, rather than progressively targeting the N-terminal (55) or C-terminal regions (52) of aSyn, it directly cleaves the central NAC and preNAC regions involved in aSyn aggregation. Moreover, no conserved active-site motif has yet been defined for TTR (56). Another major cleavage site for TTR in aSyn involves Ala^53^, which, in familial forms of PD is mutated for threonine or glutamate residues (33,57). The redox regulation of the metalloproteinase activity of TTR is possible through the S-glutathionylation of the Cys-10 residue (2,39). Indeed, we found that aSyn proteolysis by S-GSH TTR is very slow, although the presence of modifiers of aSyn conformation significantly accelerates it. Arguably, a first line of defence against misfolded extracellular aSyn might be provided by TTR, whereas MMPs and other matrix proteases are activated depending on upstream mechanisms of redox regulation.

The link between protein misfolding and the pathogenesis of PD, AD or TTR-related amyloidosis remains poorly defined especially in the non-hereditary forms of such disorders. Dramatic changes in the folding landscape of amyloidogenic proteins are induced not only by pathogenic point mutations but also by post-translational modifications and covalent bonds involving abnormal metabolites (58,59). Our study additionally shows that the proteolytic activity of TTR against aSyn is determined by conformational changes arising from physical interactions of aSyn with solid surfaces. On the side of TTR, S-glutathionylation is shown to influence the proteolytic activity of TTR, probably due to the introduction of steric hindrances to substrate binding. We describe how aSyn proteolysis is greatly enhanced by the presence of oligomeric species of TTR. Because the oxidation of the key residue Cys-10 also leads to the formation of TTR oligomers (60,61), it can be reasoned that oxidative stress accelerates aSyn clearance both directly, through the activation of MMPs, and indirectly, by sequentially promoting TTR oligomerization, aSyn misfolding and, lastly, aSyn proteolysis by tTTR. In this respect, it is relevant that the proteolytic activity of TTR targets two key regions for aSyn aggregation: the hydrophobic NAC region necessary for the aggregation onset (16) and the preNAC region that encompasses six of the seven early-onset familial PD mutations (34). Our findings warrant further research to assess the relevance of conformation-selective proteolysis of misfolded proteins *in vivo* and, more specifically, the importance of TTR-mediated degradation of aSyn in the pathogenesis of PD and ATTR-PN diseases.

## Materials and Methods

### Proteins and peptides

Different TTR preparations were used according to the protocols adopted for protein expression and purification (Table 1). In the case of preparations TTR_1st_ and Y78F_1st_, the protein was isolated by fractional ammonium sulfate precipitation as previously described (40). Briefly, recombinant TTR_1st_ and Y78F_1st_ TTR were expressed from phTTR.wt-I and phTTR.Y78F-I plasmids, respectively, based on a pET38b(+) vector (Novagene, Merck Biosciences, Darmstadt, Germany) (40,62). Both TTR_1st_ and Y78F_1st_ were expressed in *E. coli* BL21(DE3) cells using a fed-batch strategy in a 10 L bioreactor (40). The fed-batch lasted 4.5 h after which the cells were harvested and lysed. After centrifugation, the supernatant was subjected to fractional ammonium sulfate precipitation. The precipitate was dissolved in Tris buffer and dialyzed against the same buffer. The proteins were purified by anion exchange chromatography with Q-Sepharose at pH 7.6, and then dialyzed and lyophilized. The lyophilized TTR_1st_ was dissolved in 50 mM sodium phosphate pH 7.5 buffer containing 200 mM NaCl, loaded into a Superdex® 75 10/30 GL gel filtration column and the fractions corresponding to tetrameric (tTTR) and oligomeric (oTTR) TTR were collected and concentrated. In the case of preparations S-GSH-TTR_2nd_ and unmod-TTR_2nd_, the protein was isolated by preparative gel electrophoresis as previously described (3). Briefly, recombinant TTR was expressed in *E. coli* BL21 and isolated by preparative gel electrophoresis after ion-exchange chromatography. The S-GSH-TTR_2nd_ and unmod-TTR_2nd_ preparations exhibited different S-glutathionylation levels as measured by MALDI-ToF-MS: the S-GSH-TTR_2nd_ batch showed high S-GSH levels, while the unmod-TTR_2nd_ batch showed low S-GSH levels. The concentration of TTR was determined spectrophotometrically at 280 nm using calculated molar extinction coefficient values of 17780 M^−1^cm^−1^ for wild-type TTR and 16500 M^−1^cm^−1^ for Y78F. The (low) protein concentrations in samples containing only oTTR could not be quantified spectrophotometrically. Instead, equivalent TTR concentrations are estimated from the number of times the eluted sample is concentrated. Recombinant aSyn was expressed and purified as described in SI Appendix. The concentration of aSyn was determined using a Direct Detect spectrometer (Merck Millipore), an infrared-based quantification system. Depsi Aβ (1–42) peptide (click peptide) (Genscript, ref. RP10017) was dissolved in 0.1% TFA to a final concentration of 200 μM (stock solution) and stored at −80 °C until use. The peptide concentration was confirmed by UV absorption measurements at 280 nm using a Tyr extinction coefficient of 1490 M^−1^cm^−1^. Before experiments, the depsi-Aβ (1–42) peptide was thawed and converted into Aβ(1-42) by changing from acidic pH (0.1%TFA, pH 2.5) to pH > 7.4. For this, the stock solution of depsi Aβ (1– 42) peptide was diluted 1:1 using 200 mM phosphate buffer pH 7.5.

**Table 1.**
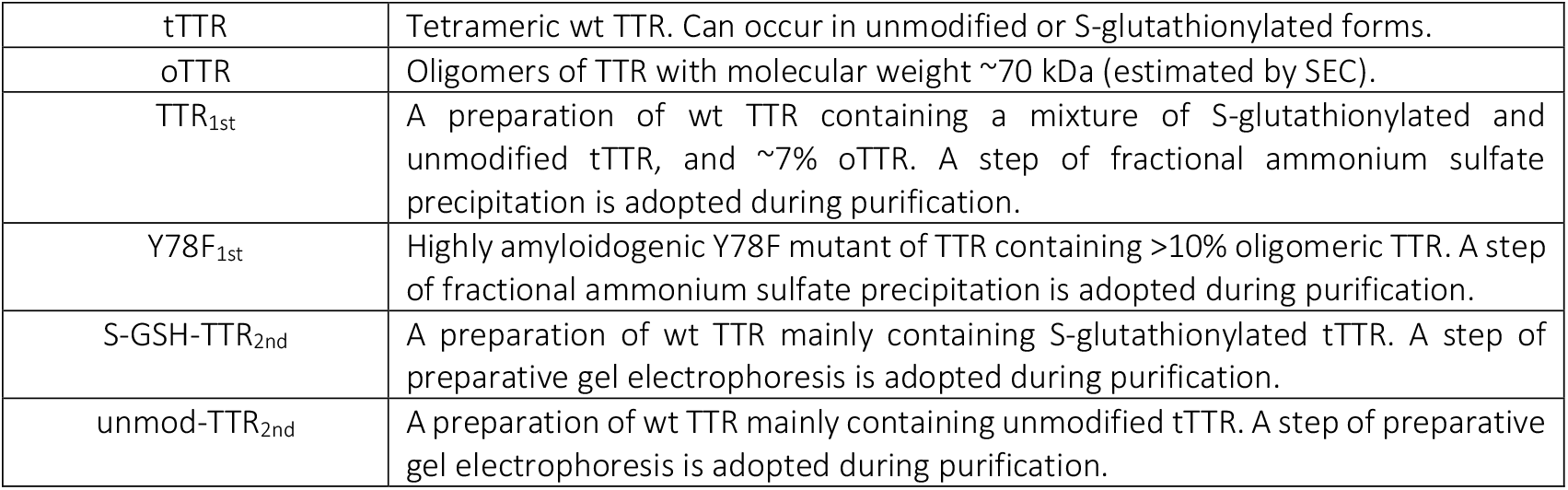
Different nomenclature and TTR preparations used in this study.

### Characterization of tetrameric and oligomeric TTR by SDS-PAGE and native immunoassays

The SEC fractions corresponding to tTTR and oTTR were collected after injection of 50 μM TTR_1st_, applied onto a 15% SDS-PAGE gel, and visualized by BlueSafe protein stain (NZYTech, ref. MB15201). To further characterize the oligomeric content of TTR_1st_, different amounts of TTR_1st_ in the range 0.1 1−1.10 μg were loaded in a native 4-15% PAGE gel (Mini-PROTEAN® TGX™ Precast Gel, Bio-Rad). The gel contained colored molecular weight standards (Precision Plus Protein Dual Color Standards, Bio-Rad), along which the gel was vertically cut. The left-side part of the gel as transferred onto a PVDF membrane (Bio-Rad) using a Trans-Blot® Turbo™ (Bio-Rad) transfer system. The membrane was immunoprobed with the polyclonal anti-TTR antibody (DAKO, ref. A0002, dilution 1:1000) and then incubated with alkaline phosphatase-conjugated polyclonal anti rabbit secondary antibody. Detection was performed with Amersham ECL® PrimeWestern blotting chemiluminescent detection reagent (Cityva) and the membrane was scanned with a ChemiDoc XRS+ imaging system (Bio-Rad) using the ImageLab 6.1 software (Bio-Rad). The right-side part of the native gel, which contained a 1.1 μg replica of TTR_1st_ and native molecular-weight standards (NativeMark™ Unstained Protein Standard, Invitrogen), was stained with BlueSafe protein stain (NZYTech).

### α-Synuclein (aSyn) aggregation followed by thioflavin-T (ThT) fluorescence

ThT fluorescence at 480 nm (440 nm excitation) was measured during the seeded aggregation of aSyn into amyloid fibrils. For this, a CHAMELEON V plate reader (HIDEX) and 384-well microplates (low flange, black, flat bottom, polystyrene; Corning, Kennebunk, ME) were used. Briefly, to serially diluted concentrations of TTR in the aggregation buffer (50 mM sodium phosphate pH 7.5 and 200 mM NaCl) was added aSyn (final concentration 2 μM), pre-formed fibrils (PFFs) of aSyn (final PFFs concentration 0.2 μM) and ThT (final concentration 70 μM) to a final volume of 50 μL. The microplate was incubated at 37°C and ThT fluorescence was measured every 30 min for approximately 180 h. To prevent evaporation, 30 μL paraffin oil was added to each reaction well with an automatic multichannel pipette (Eppendorf Xplorer, ref 4861000120). Average fluorescent values were computed for four independent experiments run for each TTR concentration, or for twelve independent experiments run in the absence of TTR. The aSyn PFFs were produced by incubating 500 μL samples of 357 μM aSyn on a Thermo shaker incubator (MSC-100 Cooling Thermoshaker Incubator, Labgene Scientific) at 37°C and 1000 rpm. After 8 days of incubation, the samples were diluted in aggregation buffer to a final concentration of 50 μM aSyn and sonicated for 2 min (SONOPULS HD2200, Bandelin Electronic, probe: MS73) during on/off sequences of 15 seconds each (10% max power). The aSyn PFFs thus produced were frozen in liquid nitrogen and kept at −80 °C.

### Western blot and kinetic analyses of aSyn and Aβ Proteolysis by TTR

Unless specified otherwise, all proteolysis experiments were carried out in 1.5 mL Protein LoBind polypropylene tubes (Eppendorf, ref.0030108.116), in the aggregation buffer and under quiescent conditions at 37 °C. The effect of different TTR preparations (5 μM TTR) on the proteolysis of 8 μM aSyn was investigated in the absence and presence of a cylindrical 10 mm×3 mm Teflon stirring bar (Merck, ref. Z328650) or three 4.5 mm diameter glass beads (Zymo Research, ref. S1001). Aβ(1-42) proteolysis by TTR was also studied by incubating 20 μM Aβ(1-42) with 10 μM TTR_1st_. In all cases, aliquots of the reaction mixtures were collected at different timepoints, run on a 15% or 20% SDS-PAGE gel, and transferred onto a nitrocellulose membrane (Bio-Rad) using a Trans-Blot® Turbo™ (Bio-Rad) transfer system. Monoclonal anti-aSyn antibody 3C11 (StressMarq, ref.SMC-530, 1:1000 dilution) or monoclonal anti-Aβ antibody BAM-10 (Sigma, ref. A3981, 1:2000 dilution) were used for the immunodetection of aSyn or Aβ(1-42), respectively. In both cases, alkaline phosphatase-conjugated polyclonal anti-mouse antibody (Sigma, 1:10,000 dilution) was used as secondary antibody. Detection was performed with Amersham ECL® PrimeWestern blotting chemiluminescent detection reagent (Cityva) and the membranes were scanned with a ChemiDoc XRS+ imaging system (Bio-Rad) using the ImageLab 6.1 software (Bio-Rad). The density decay was expressed in normalized units and used to estimate specificity rate constants (k_cat_/K_M_) assuming pseudo-first-order conditions (2).

### Matrix-Assisted Laser Desorption/Ionization coupled to Time-of-Flight Mass Spectrometry (MALDI-ToF-MS)

A saturated solution of sinapinic acid (SA) as matrix in 30:70 (v/v) acetonitrile:water, 0.1% trifluoroacetic acid (TFA) and a protein solution in 30:70 (v/v) acetonitrile:water, 0.1% TFA were prepared and mixed in a 1:1 ratio. A total of 1 μL of the previous mixture was deposited into a polished stainless-steel target (Bruker, Bremen, Germany) and allowed to dry. Then, 1 μL of SA matrix solution was deposited into the sample and allowed to dry. The same procedure was followed for the Protein Standard Calibration I solution (Bruker) used for Calibration. The target was introduced in a Autoflex MALDI-TOF (Bruker), spectra were acquired in lineal mode (Flex control, Bruker) and processed by Flex Analysis (Bruker).

### nanoLiquid Chromatography-Mass Spectrometry (nanoLC-MS/MS)

Samples were diluted to a final 0.5 μM concentration in a mixture of 10% acetonitrile (ACN), 1% formic acid (FA) aqueous solution for nanoLC-MS/MS. The analysis of aSyn (P37840) and TTR (P02766) protein mixtures was conducted on a NanoAcquity Ultra Performance chromatographic system (Waters Corp., Milford, MA, US) coupled to an Orbitrap Fusion Lumos tribrid mass spectrometer (Thermo Scientific). We used the Advion TriVersa NanoMate (Advion) as the nanospray interface through chip technology. Samples were automatically injected to the chromatographic system through the Waters Sample Manager into a C4 trap column (180 μm i.d. × 20 mm, 5 μm, 300 Å, Acquity UPLC M-Class Symmetry, Waters). Protein content was eluted and separated using an Acquity UPLC M-Class BioResolve mAb column (75 μm × 250 mm, 2.7 μm, 450 Å) (Waters) with linear consecutive gradients from 10 to 50 % B in 60 min and from 50% to 85% in 7min, at a flow rate of 300nl/min (A= 0.1 % FA in water, B= 0.1 % FA in ACN). The mass spectrometer was operated at low pressure (2 mbar), in data-dependent acquisition (DDA) mode and positive polarity. Survey MS scans were acquired in the Orbitrap with the resolution (defined at 200 m/z) set to 120,000. The top speed (most intense) ions per scan were fragmented and detected in the Orbitrap with the resolution set to 60,000. We used different acquisition methods with two types of fragmentation techniques: a) HCD (High collision dissociation) and b) EThcD (electron-transfer dissociation with supplemental high collision dissociation). The ion count target value was 400,000 and 500,000 for the survey scan and the MS/MS scan, respectively in all methods. Target ions already selected for MS/MS were dynamically excluded for 30 s. RF Lens were tuned to 30 %. For the HCD method, we used a 20% collision energy, 300 ms maximum injection time and included charge states 4 to 40 for fragmentation with minimum intensities thresholds required to trigger MS to MS/MS switch at 25000. For the EThcD, we differentiated charges 4 to 10 and 7 to 24 for fragmentation, using 25 ms and 10 ms of ETD reaction time respectively. The ETD reagent target was set to 700000 and 200 ms and supplemental HCD energy to 10%. Maximun injection time was set to 300 ms. The analysis of the nanoLC-MS/MS data was performed as described in SI Appendix.

### Infrared fluorescent western blot analysis of aSyn conformations and oligomeric states

To study the effects of Teflon surfaces and TTR oligomers on aSyn conformation, samples of 8 μM aSyn in the aggregation buffer were incubated at 37 °C under quiescent conditions and over 20 h with and without a cylindrical 10 mm x 3mm Teflon stirring bar (Merck, ref. Z328650) or 0.5 μM oTTR. Aliquots of the reaction mixtures were collected, run in a native 4-15% PAGE gel (Mini-PROTEAN® TGX™ Precast Gel, Bio-Rad), and then transferred onto a nitrocellulose (Bio-Rad) membrane using a Trans-Blot® Turbo™ transfer (Bio-Rad) system. Possible ASyn conformers or oligomers on the membrane were probed using monoclonal anti-aSyn antibody 3C11 (StressMarq, ref.SMC-530, 1:1000 dilution). For high sensitivity and low background signal, infrared fluorophore conjugated anti-mouse antibody IRDye 800CW (Li-COR, ref. 926-32210, dilution 1:20,000) was used as secondary antibody. Direct infrared fluorescence detection was performed using an Odyssey CLx Infrared Imaging System (Li-COR) and the Image Studio™ (Li-COR) signal identification software. The images were analysed by Empiria Studio (LI-COR) data integrity software.

### Dynamic Light Scattering (DLS)

The formation of aSyn aggregates in the presence of TTR_1st_ was investigated using DLS. For that, 200 μL samples of 4 μM aSyn containing 1 μM TTR_1st_ were incubated in a micro UV-cuvette with cap (BRAND GMBH, ref.759200) at 37 °C under quiescent over 48 h. DLS was also employed to monitor the formation of aSyn aggregates in the absence of TTR. In this case, 380 μL samples of 8 μM aSyn solutions were incubated in Protein LoBind polypropylene tubes (Eppendorf, ref.0030108.116) at 37 °C under quiescent conditions over 10 days in the absence and presence of a cylindrical 10 mm x 3mm Teflon stirring bar (Merck, ref. Z328650) or three 4.5 mm diameter glass beads (Zymo Research, ref. S1001). Aliquots (70 μL) of the reaction mixtures were collected at different timepoints and analysed in a micro UV-cuvette with cap (BRAND GMBH, ref.759200) using a Zetasizer Nano ZS DLS system (Malvern Instruments) at a scattering angle of 173°. The Zetasizer Nano software (Malvern Instruments) was used to determine Intensity-based size distributions and the Derived Count Rate (an indicator of the overall scattering intensity).

### Transmission Electron Microscopy (TEM) analysis of aSyn fibrils and oligomers

The species produced during the seeded and unseeded aggregation of aSyn were visualized using a JEM-1400 (JEOL, Tokyo, Japan) TEM at an accelerating voltage of 80 kV. For that, the collected aliquots were adsorbed onto glow-discharged, carbon-coated films supported on 300-mesh nickel grids and negatively stained with 1% (w/v) uranyl acetate.

## Acknowledgements

This work is part of a project that has received funding from the European Union’s Horizon 2020 research and innovation programme under grant agreement No. 952334 (PhasAGE). This research was funded by the Portuguese Foundation for Science and Technology (FCT) in the framework of projects PTDC/QUI-COL/2444/2021 and PTDC/MED-PAT/0959/2021. T.G. is a recipient of a fellowship by FCT (ref. 2020.07444.BD). M.A.L. is supported by CEECINST/00091/2018. A.P. acknowledges support from Grant PID2019-104350RB-I00 and G.A. acknowledges support from Grant PID2020-117656RB-I00, both grants from the Ministry of Science and Innovation (MICINN), Spain.

We thank Dr. Marta Vilaseca, Institute for Research in Biomedicine (IRB Barcelona), for the nanoLC-MS/MS analysis and Patricia Torruella, Laboratory of Biochemistry, IQS, University Ramon Llull, Barcelona for the MALDI-TOF experiments.

