## Supporting Information for "Transthyretin has conformation-selective proteolytic activity against α-synuclein"

### Supplementary Methods

**α-Synuclein (aSyn) expression and purification.** Human aSyn was expressed and purified by adapting a protocol from Pujols et al. (1). Accordingly, the cDNA of aSyn in pT7-7 plasmid (Addgene, Watertown, USA) was transformed and overexpressed in Escherichia coli BL21 (DE3) cells in LB medium containing 100 µM/mL ampicillin. Protein expression at 37 °C was induced with 1 mM IPTG when OD600 reached 0.6. Cells were harvested after 4 h of expression by centrifugation (4000 xg, 20 min at 4 °C) and frozen at –80 °C. The thawed pellet was lysed on ice by sonication (SONOPULS HD2200, Bandelin Electronic) in lysis buffer (50 mM Tris-HCl pH 8, 150 mM NaCl, 1 µg/mL pepstatin, 20 µg/mL aprotinin, 1 mM benzamidine, 1mM phenylmethylsulfonyl fluoride (PMSF), 1 mM EDTA and 0.25 mg/mL lysozyme). The obtained cell extract was boiled at 95 °C for 10 min and clarified by centrifugation (17000 xg, 45 min at 4 °C). Then 136 µL/mL of 10% w/v streptomycin sulfate and 228 µL/mL of pure acid acetic were added to the supernatant and centrifuged (17000 xg, 10 min at 4 °C). The aSyn in the soluble fraction was precipitated by saturated ammonium sulfate (550 g/L) 1:1 (v/v) and centrifuged (17000 xg, 10 min at 4 °C). The pellet was washed first with 50% ammonium sulfate, then with 100 mM ammonium acetate pH 8.0 (5 mL/L culture) and, finally with pure EtOH 1:1 (v/v). After each washing step, the pellet was separated by centrifugation (17000 xg, 10 min at 4 °C). The final pellet was resuspended, filtered (0.45 µm filter) and buffer exchanged to 20 mM Tris-HCl pH 8.0 using a HiPrep26/10 desalting column (GE Healthcare). The obtained soluble aSyn was loaded on a Resource-Q anion exchange column (6 mL, GE Healthcare) and eluted with 20-column volume linear gradient of NaCl (0-500 mM). Fractions containing pure aSyn were pooled, concentrated (10 kDa cutoff ultrafiltration device, Merck Millipore) and further purified by size exclusion chromatography using a HiPrep26/60 Sephacryl S-100 column (GE Healthcare) equilibrated with protein buffer (50 mM sodium phosphate pH 7.5, 200 mM NaCl). Eluted fractions containing pure aSyn (analysed by 15% SDS-PAGE) were pooled and concentrated. The pure aSyn was frozen in liquid nitrogen and kept at –80 °C.

**Analysis of nanoLiquid Chromatography-Mass Spectrometry (nanoLC-MS/MS) data.** To perform the MS data analysis for proteoform identification and relative quantitation at 3 time points (15 min, 30 min and 45 min) two different software/search nodes were employed: ProSight PD and BioPharma Finder. Proteome Discoverer v2.5.0.400 (Thermo Fisher Scientific) with ProSightPD node (2) was used for proteoform identification. The software assigns MS1 and the corresponding MS2 spectra to protein sequences in the database. Database searches in ProSightPD were performed using an xml-annotated database from SwissProt (Human-release 2021/04, containing 16676 entries) translated to a .psdb format through the ProSightPD database tool inside Proteome Discoverer software. A 3-tier iterative search (Absolute, Biomarker and Absolute with delta M mode) was performed with precursor mass tolerances of 2.2 Da / 10 ppm /25000 Da respectively and fragment mass tolerances of 10 ppm. The iterative search approach allows for a reduction in data processing time by short-circuiting further, more computationally intensive searches once a statistically confident identification is made. Acetylation in protein N-terminus was introduced as a dynamic modification. Proteoform spectrum matches with FDR < 1% were considered as positive hits. ProSight Lite (3), integrated in Prosight PD node was used for further manual proteoform validation. BioPharma Finder v.4 (Thermo Fisher Scientific) was used to extract averaged mass spectra at the MS1 level from the total ion chromatogram and to generate a list of deconvoluted masses with corresponding ion intensities (Fig. 3). Deconvolution to the monoisotopic zero charged proteoform masses was done using auto Xtract algorithm on resolved m/z charged species (4,5). Deconvoluted masses were manually assigned to aSyn (P37840) and TTR (hTTR; P02766) proteoforms by match between the experimental and theoretical masses and by concordance with the proteoform identification based on the M1 and MS2 spectra derived from ProSightPD database searches.

### Figures S1 to S8

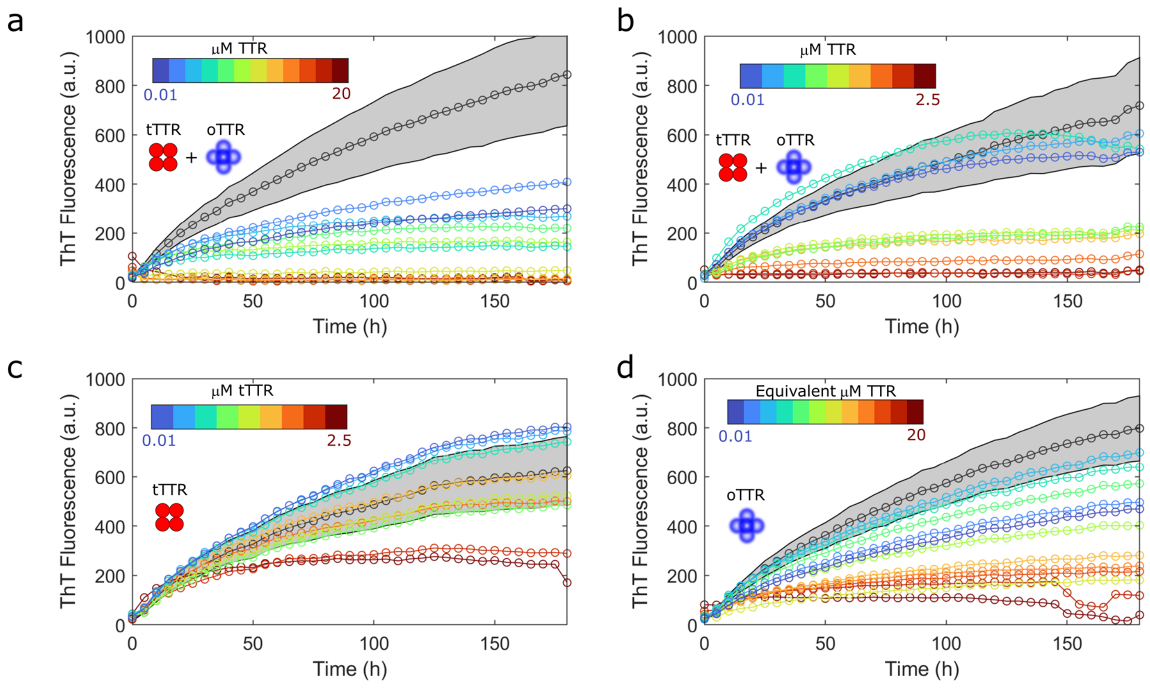

Figure S1. Effect of TTR on the aggregation progress curves of 2 μM aSyn measured in the presence of 0.2 μM PFFs. (a to d) Dose-response studies using TTR solutions serially diluted from (a) 20 μM TTR, (b) 2.5 μM TTR, (c) 2.5 μM tTTR and (d) oTTR solutions concentrated to the equivalent of 20 μM tTTR. Color bars: sequence of serial dilutions from the starting TTR concentration (dark red) to 0.01 μM TTR (dark blue). Grey symbols and shaded areas represent averages and standard deviations of ThT fluorescence measured for control experiments without added TTR. (a and b) The proteolytically-active $TTR_{1st}$ preparation (Table 1) was used. (c and d) The samples (c) tTTR and (d) oTTR derive from the $TTR_{1st}$ preparation.

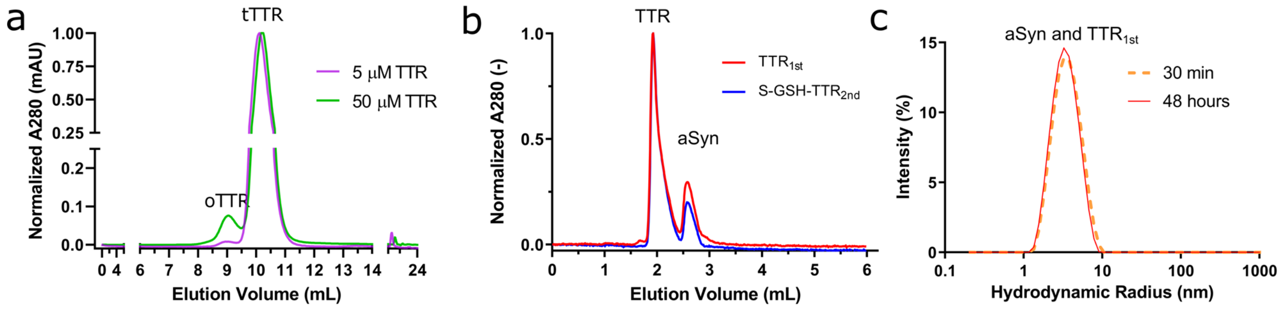

Figure S2. Further characterization of the protein samples. (a) The amount of oTTR changes with the total concentration of $TTR_{1st}$. Normalized SEC chromatograms of 5 and 50 μM TTR (Superdex® 75 10/30 GL column). (b and c) α-Synuclein proteolysis produces no detectable aggregates. (b) Normalized SEC chromatograms of 2 μM aSyn incubated over 3 days with 1 μM TTR (Superdex® 200 Increase 5/150 GL column). Proteolytically-active ($TTR_{1st}$) and inactive (S-GSH-$TTR_{2nd}$) preparations were tested. (c) DLS analysis of 4 μM aSyn incubated with 1 μM $TTR_{1st}$ over the indicated time intervals.

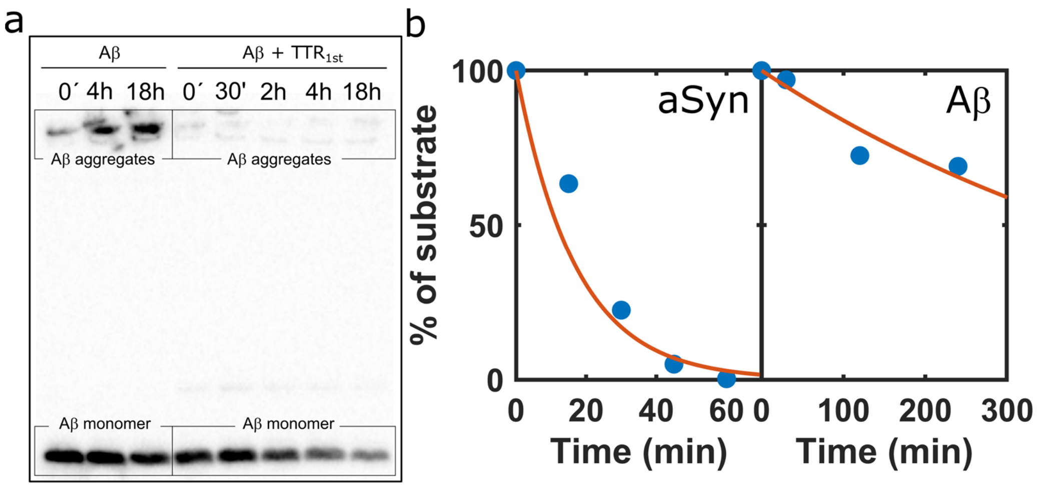

Figure S3. Comparison of the proteolytic activity of $TTR_{1st}$ using aSyn or Aβ-peptide substrates. (a) Western blot analysis of aliquots of 20 μM Aβ incubated with 10 μM $TTR_{1st}$ over the indicated time intervals. The aliquots were immunoblotted with the anti-Aβ antibody. (b) Symbols: fractional decay of aSyn (left) and Aβ (right) with time estimated by quantitative image analysis of the immunoblotted aliquots in Figs. 2b and S3a, respectively. Lines: numerical fittings by a pseudo-first-order progress curve (aSyn: $k_{cat}/K_M$=197 $s^{-1}M^{-1}$; Aβ: $k_{cat}/K_M$=2.9 $s^{-1}M^{-1}$).

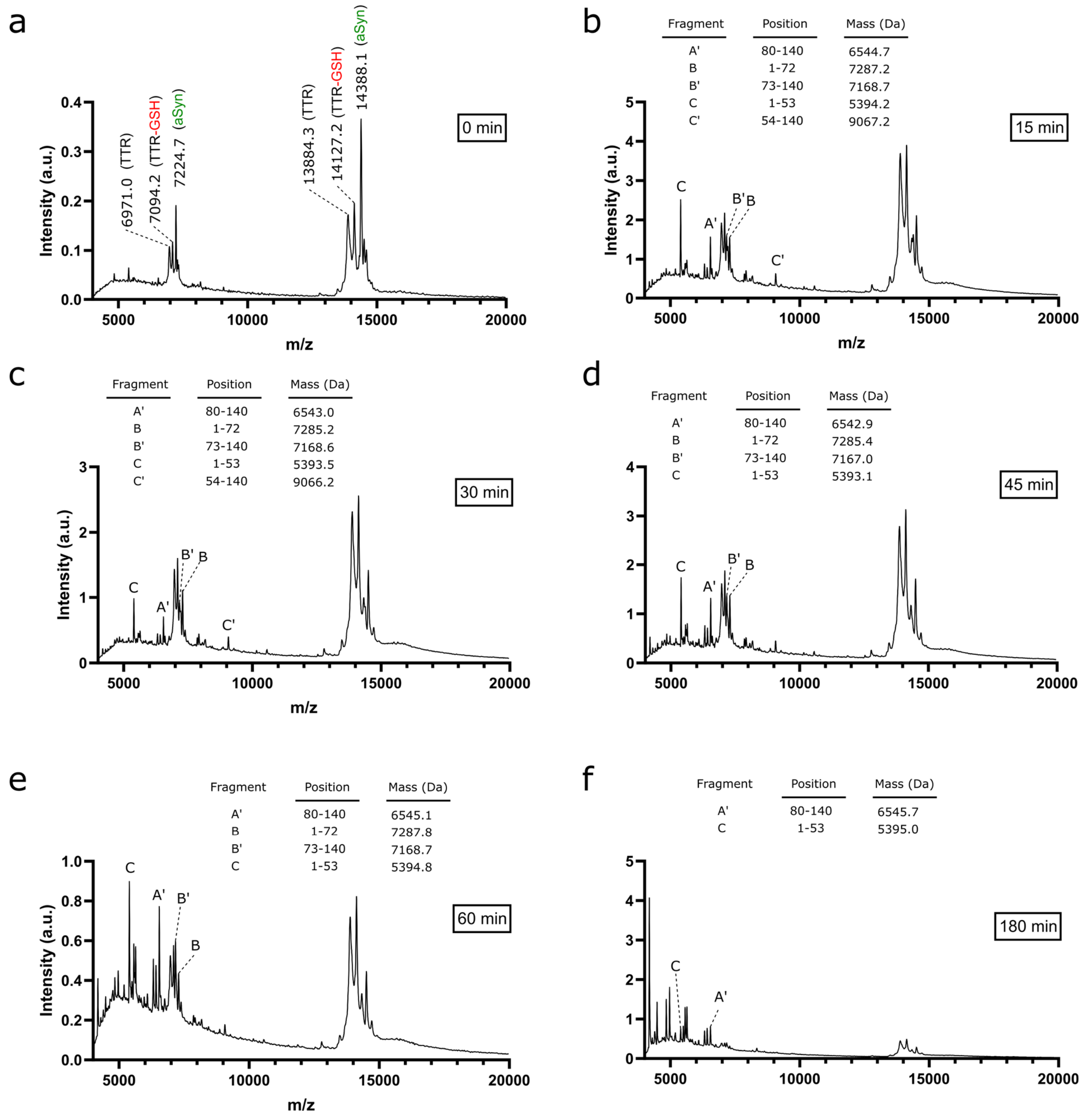

Figure S4. MALDI-ToF-MS monitoring of aSyn (8 μM) proteolysis by TTR$_{1st}$ (5 μM). Spectra of (a) freshly prepared solutions, or (b to f) incubated over (b) 15, (c) 30, (d) 45, (e) 60, and (f) 180 min. (a) Next to the peaks are given the molecular mass (*m/z,* mass-to-charge values) of aSyn, unmodified TTR, and S-GSH TTR. For clarity, this information is not provided in the subsequent panels. (b to f) Letters next to the peaks indicate different aSyn fragments following the same notation as in Fig. 3. Tables: start/stop residue positions and observed mass of each fragment.

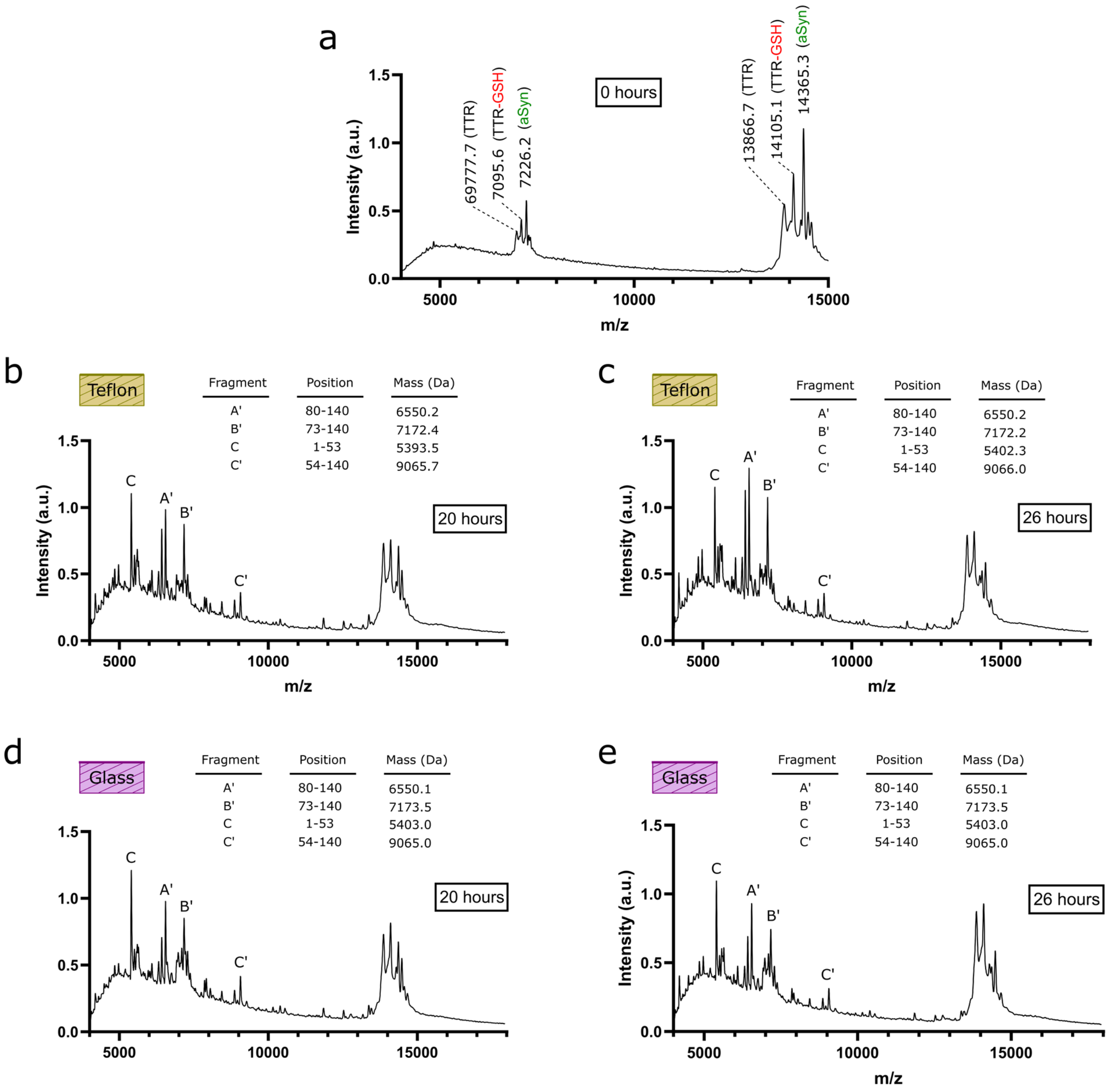

Figure S5. MALDI-ToF-MS monitoring of aSyn (8 μM) proteolysis by tTTR (5 μM) in the presence of different surfaces. Spectra of (a) freshly prepared solutions, or (b and c) incubated in the presence of Teflon surfaces over (b) 20, and (c) 26 h, or (d and e) glass surfaces over (d) 20, and (e) 26 h. (a) Next to the peaks are given the molecular mass (*m/z,* mass-to-charge values) of aSyn, unmodified TTR, and S-GSH TTR. For clarity, this information is not provided in the subsequent panels. (b to e) Letters next to the peaks indicate different aSyn fragments following the same notation as in Fig. 3. Tables: start/stop residue positions and observed mass of each fragment.

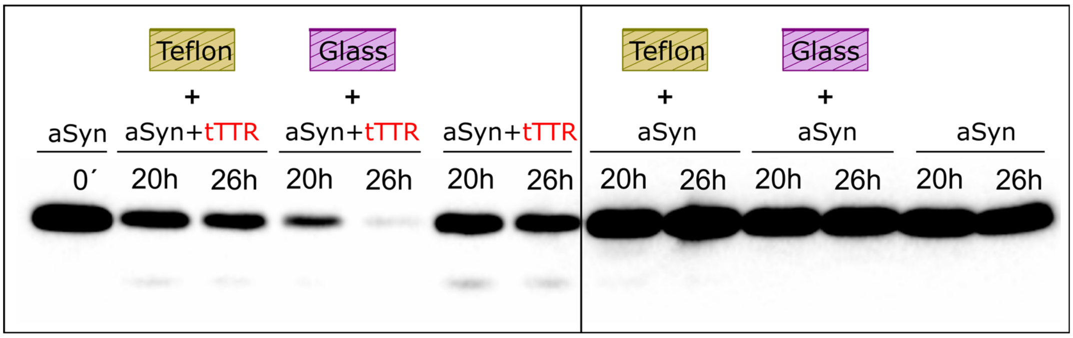

Figure S6. The effect of Teflon and glass surfaces on the proteolysis of aSyn by tTTR – additional data. Western blot analysis of 8 μM aSyn aliquots incubated with (left panel) and without (right panel) tTTR in the presence of Teflon and glass surfaces over the indicated time intervals. Left panel: same as Fig. 4a. Right panel: Control experiments showing that the results in the left panel are not explained by protein depletion caused by aSyn aggregation.

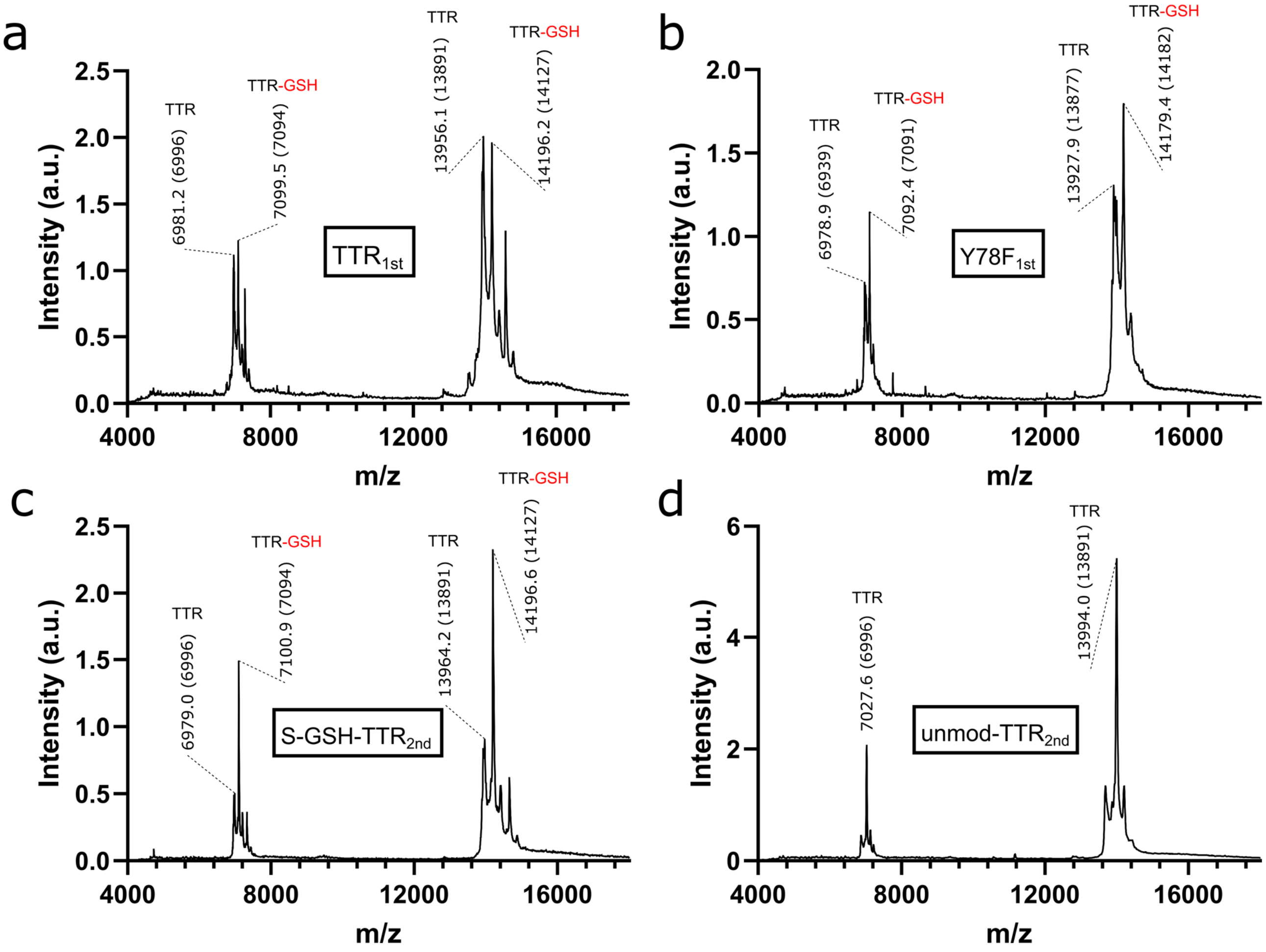

Figure S7. MALDI-ToF-MS spectra of the different TTR preparations used in this study. The protein production protocol of (a) $TTR_{1st}$ and (b) $Y78F_{1st}$ TTR comprises a step of fractional ammonium sulfate precipitation, while the preparation of S-GSH-$TTR_{2nd}$ and unmod-$TTR_{2nd}$ comprises a step of preparative gel electrophoresis. Next to the peaks are given the measured and (in parenthesis) theoretical values of the molecular mass (*m/z*) of unmodified and S-GSH TTR.

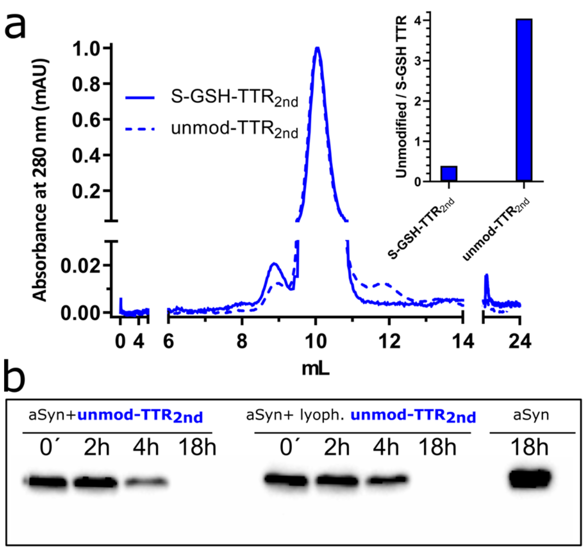

Figure S8. S-glutathionylation of TTR inhibits its proteolytic activity against aSyn. (a) Normalized SEC chromatograms of S-glutathionylated (S-GSH-$TTR_{2nd}$, solid line) and unmodified (unmod-$TTR_{2nd}$, dashed line) TTR preparations produced by the same expression and purification protocol. Inset: ratios of unmodified to S-GSH TTR estimated from the MALDI-ToF-MS spectra of “inactive” S-GSH-$TTR_{2nd}$ (Fig. S7c) and “active” unmod-$TTR_{2nd}$ (Fig. S7d). (b) Western blot analysis of aSyn aliquots incubated with unmod-$TTR_{2nd}$ (left series) over the indicated time intervals. The lyophilization of the unmod-$TTR_{2nd}$ sample does not affect its proteolytic activity significantly (middle series). Far-right lane: negative control.

### Table S1

**Table S1.** Complementary data to Fig. 3. Overall proteoforms identified by Top-down MS using ProsightPD. Columns depict: a) Experimental monoisotopic deconvoluted masses (in Da), b) Uniprot Accession Number, c) Start and d) End amino acid positions in the original protein sequence, e) Identified proteoform sequence f) MS fragmentation method used to acquire the MS2 spectra, g),h),i) MS peak intensity at 15, 30 and 45 min, respectively.

| Mass (Da) | Protein | AA start | AA End | Proteoform sequence | Frag type | Intensity 15min | Intensity 30min | Intensity 45min |
|---|---|---|---|---|---|---|---|---|
| 14451.231 | P37840 · SYUA_HUMAN | 1 | 140 | MDVFMKGLSKAKEGVVAAAEKTKQGVAEAAGKTKEGVLYVGSKTKEG VVHGVATVAEKTKEQVTNVGGAVVTGVTAVAQKTVEGAGSIAAATGF VKKDQLGKNEEGAPQEGILEDMPVDPDNEAYEMPSEEGYQDYEPEA | HCD | 43748340 | 38898371 | 8613289 |
| 4171.681 | P37840 · SYUA_HUMAN | 104 | 140 | EEGAPQEGILEDMPVDPDNEAYEMPSEEGYQDYEPEA | EThcD | 1880388 | 4918434 | 13500089 |
| 5387.889 | P37840 · SYUA_HUMAN | 1 | 53 | MDVFMKGLSKAKEGVVAAAEKTKQGVAEAAGKTKEGVLYVGSKTKEG VVHGVA | EThcD | 9510102 | 76832 | 14089512 |
| 5558.442 | P37840 · SYUA_HUMAN | 91 | 140 | ATGFVKKDQLGKNEEGAPQEGILEDMPVDPDNEAYEMPSEEGYQDYE PEA | HCD | 997262 | 3250617 | 203688 |
| 5629.480 | P37840 · SYUA_HUMAN | 90 | 140 | AATGFVKKDQLGKNEEGAPQEGILEDMPVDPDNEAYEMPSEEGYQDY EPEA | HCD | 1241489 | 3740723 | 250965 |
| 6542.968 | P37840 · SYUA_HUMAN | 80 | 140 | KTVEGAGSIAAATGFVKKDQLGKNEEGAPQEGILEDMPVDPDNEAYE MPSEEGYQDYEPEA | HCD | 4316337 | 13201426 | 1005629 |
| 6602.539 | P37840 · SYUA_HUMAN | 1 | 64 | MDVFMKGLSKAKEGVVAAAEKTKQGVAEAAGKTKEGVLYVGSKTKEG VVHGVATVAEKTKEQVT | EThcD | 5165774 | 5632752 | 7242055 |
| 7169.310 | P37840 · SYUA_HUMAN | 73 | 140 | GVTAVAQKTVEGAGSIAAATGFVKKDQLGKNEEGAPQEGILEDMPVDP DNEAYEMPSEEGYQDYEPEA | HCD | 4567708 | 13075483 | 830188 |
| 7169.310 | P37840 · SYUA_HUMAN | 73 | 140 | GVTAVAQKTVEGAGSIAAATGFVKKDQLGKNEEGAPQEGILEDMPVDP DNEAYEMPSEEGYQDYEPEA | EThcD | 25818616 | 21178093 | 27307874 |
| 7299.914 | P37840 · SYUA_HUMAN | 1 | 72 | MDVFMKGLSKAKEGVVAAAEKTKQGVAEAAGKTKEGVLYVGSKTKEG VVHGVATVAEKTKEQVTNVGGAVVT | EThcD | 10975219 | 10109406 | 14537165 |
| 7926.258 | P37840 · SYUA_HUMAN | 1 | 79 | MDVFMKGLSKAKEGVVAAAEKTKQGVAEAAGKTKEGVLYVGSKTKEG VVHGVATVAEKTKEQVTNVGGAVVTGVTAVAQ | HCD | 1209527 | 5166142 | 0 |
| 9081.331 | P37840 · SYUA_HUMAN | 54 | 140 | TVAEKTKEQVTNVGGAVVTGVTAVAQKTVEGAGSIAAATGFVKKDQLG KNEEGAPQEGILEDMPVDPDNEAYEMPSEEGYQDYEPEA | HCD | 2124909.6 | 4744329.4 | 1664471.2 |
| 9082.862 | P37840 · SYUA_HUMAN | 1 | 92 | MDVFMKGLSKAKEGVVAAAEKTKQGVAEAAGKTKEGVLYVGSKTKEG VVHGVATVAEKTKEQVTNVGGAVVTGVTAVAQKTVEGAGSIAAAT | HCD | 847806 | 2117223.6 | 46514.68 |
| 13915.921 | P02766 · TTHY_HUMAN+ met+cystsulfinic acid (31) | 1 | 128 | MGPTGTGESKCPLMVKVLDAVRGSPAINVAVHVFRKAADDTWEPFAS GKTSESGELHGLTTEEEFVEGIYKVEIDTKSYWKALGISPFHEHAEVVFTA NDSGPRRYTIAALLSPYSYSTTAVVTNPKE | HCD | 20741959 | 183937797 | 88593392 |
| 14189.000 | P02766 · TTHY_HUMAN+ met+S-glutathione (306) | 1 | 128 | MGPTGTGESKCPLMVKVLDAVRGSPAINVAVHVFRKAADDTWEPFAS GKTSESGELHGLTTEEEFVEGIYKVEIDTKSYWKALGISPFHEHAEVVFTA NDSGPRRYTIAALLSPYSYSTTAVVTNPKE | HCD | 64777783 | 222829417 | 101120430 |
